# Deciphering GB1’s Single Mutational Landscape: Insights from MuMi Analysis

**DOI:** 10.1101/2024.06.01.596930

**Authors:** Tandac F. Guclu, Ali Rana Atilgan, Canan Atilgan

**Author notes:** **Correspondence**: Tandac Furkan Guclu, Faculty of Natural Sciences and Engineering, Sabanci University, Tuzla 34956 Istanbul, Turkey.

## Abstract

Mutational changes that affect the binding of the C2 fragment of Streptococcal protein G (GB1) to the Fc domain of human IgG (IgG-Fc) have been extensively studied using deep mutational scanning (DMS), and the binding affinity of all single mutations has been measured experimentally in the literature. To investigate the underlying molecular basis, we perform *in-silico* mutational scanning for all possible single mutations, along with 2-µs-long molecular dynamics (WT-MD) of the wild-type (WT) GB1 in both unbound and IgG-Fc bound forms. We compute the hydrogen bonds between GB1 and IgG-Fc in WT-MD to identify the dominant hydrogen bonds for binding, which we then assess in conformations produced by Mutation and Minimization (MuMi) to explain the fitness landscape of GB1 and IgG-Fc binding. Furthermore, we analyze MuMi and WT-MD to investigate the dynamics of binding, focusing on the relative solvent accessibility (RSA) of residues and the probability of residues being located at the binding interface. With these analyses, we explain the interactions between GB1 and IgG-Fc and display the structural features of binding. Our findings pave the way for improved predictive accuracy in protein stability and interaction studies, which are crucial for advancements in drug design and synthetic biology.

## INTRODUCTION

The adaptive landscape of proteins arising from point mutations refers to the extent of amino acid replacements that are allowed in a protein sequence in a given background.^1^ These mutations can result from factors such as errors in DNA replication, contact with mutagens or natural genetic variation. Understanding the fitness landscape of point mutations is vital in fields such as molecular evolution, protein engineering and personalized medicine, as different amino acid substitutions can have various effects on protein structure, stability and function.^2^ Experimentally, deep mutation scanning (DMS) is a powerful technique used in the comprehensive analysis of the functional consequences of amino acid changes.^3^ With DMS, mutations are systematically introduced at every position in the protein and the effects of these changes are measured. However, DMS can be resource intensive and costly, especially when performed on a large scale or with multiple iterations. The limitations in DMS and the potential impact of single mutations on protein function to trigger many discoveries in the biochemical sciences has driven researchers to develop the relatively inexpensive computational methods^4^ whereby algorithms to analyze and interpret large-scale data generated by DMS experiments are developed.^5^ These approaches play a critical role in extracting meaningful insights from the vast amounts of sequencing data generated by DMS. Computational tools are used to predict the functional effects of amino acid substitutions based on features such as protein sequence, structure and evolutionary conservation. These tools aim to identify possible mutations that could alter protein function, stability or interactions; see e.g.^6^ This approach includes identifying mutations that significantly alter protein function by comparisons with the wild type (WT) sequence, mapping functionally important spots, and regions of structural importance. Meanwhile, machine learning techniques such as artificial neural networks and random forests are increasingly used to analyze DMS data. These approaches learn complex patterns from the data and can make predictions about the effects of mutations on protein function. Computational methods integrate protein sequences as well as structural biology data such as protein structures and molecular dynamics (MD) simulations.^7^ Web servers and databases created as a result of these efforts provide tools and resources for analyzing DMS data; see e.g.^8^

Chief among the limitations of computational approaches is its low prediction accuracy. Although many prediction algorithms have been developed, they are often based on simplified models of protein structure and function and may not fully reflect the complexity of real biological systems.^9^ Factors such as protein-protein interactions, post-translational changes, cellular localization and genetic background can modulate the functional consequences of mutations but may not be adequately accounted for in computational models. On the other hand, the accuracy and reliability of computational predictions depend crucially on the quality of the experimental data used to train and validate the prediction models. Moreover, some of the more sensitive methods require intensive use of computational resources, imposing significant constraints on researchers without access to high-performance computing infrastructure. Therefore, there is need to develop approaches that can produce fast, yet consistent, predictions with reasonable computational resources by not requiring lengthy MD simulations while accounting for ambient conditions such as water and ionic strength.

While sufficient data for machine learning methods is provided by DMS experiments, information on structural details lags far behind.^5^ Methods that account for external effects such as water and salt are required to track the small but significant changes in protein structure caused by point mutations in sufficient detail. Foremost among these are MD simulations and related advanced simulation techniques, which require gross computational time.^10^ The use of data from MD simulations, which are computationally expensive, in combination with machine learning techniques to overcome the lack of structural information has only recently been used.^11^

The processes contributing to the cumulative effects observed in DMS experiments can be calculated in a way that decouples folding and binding processes.^9,12^ On the down side, although mutation effects have been quantified and calculated as energy differences compared to the WT protein, the underlying molecular mechanisms leading to these values have not been identified. Previous efforts to find an atomic-scale link in the mechanisms leading to changes in conformational values in DMS have shown that the solvent accessible surface is informative about mutational effects on the binding affinity of hydrogen bonds and salt bridges.^12–15^ In particular, the classification of residues based on solvent accessibility^12,13,15^ without any measurement of binding energies shows promise. However, methods to predict the stability of GB1 using these factors as attributes in machine learning algorithms have fallen short of predicting the experimental findings to the desired degree.^12,13,15^

A common method to assess the fitness is by measuring binding affinity, which has been extensively studied by deep mutational scanning (DMS) experiments. As a model system, the binding profile of all possible single and double mutations belonging to C2 fragment of Streptococcal protein G (GB1) and Fc domain of human IgG (IgG-Fc) have been measured experimentally (see Figure 1 for structures).^16^ Furthermore, several publications discussed the effects of these mutations by changing up to four positions in the sequence^17^ and by stability experiments.^18,19^ GB1 was also used as a case study to predict protein structures by only using epistatic relations between residue pairs.^20,21^ The processes contributing to the observed cumulative effects in the DMS experiments were separated into those due to binding fitness and folding fitness.^9,12^ Although the effects of mutations are measured with respect to the WT and calculated as energy differences, the underlying molecular mechanisms leading to these values are not well-established. Moreover, methods to predict stability of GB1 by using factors such as solvent accessible surface, hydrogen bonds and salt-bridges as features for machine learning algorithms are not sufficiently correlated with the experiments.^18,22^

**Figure 1.**
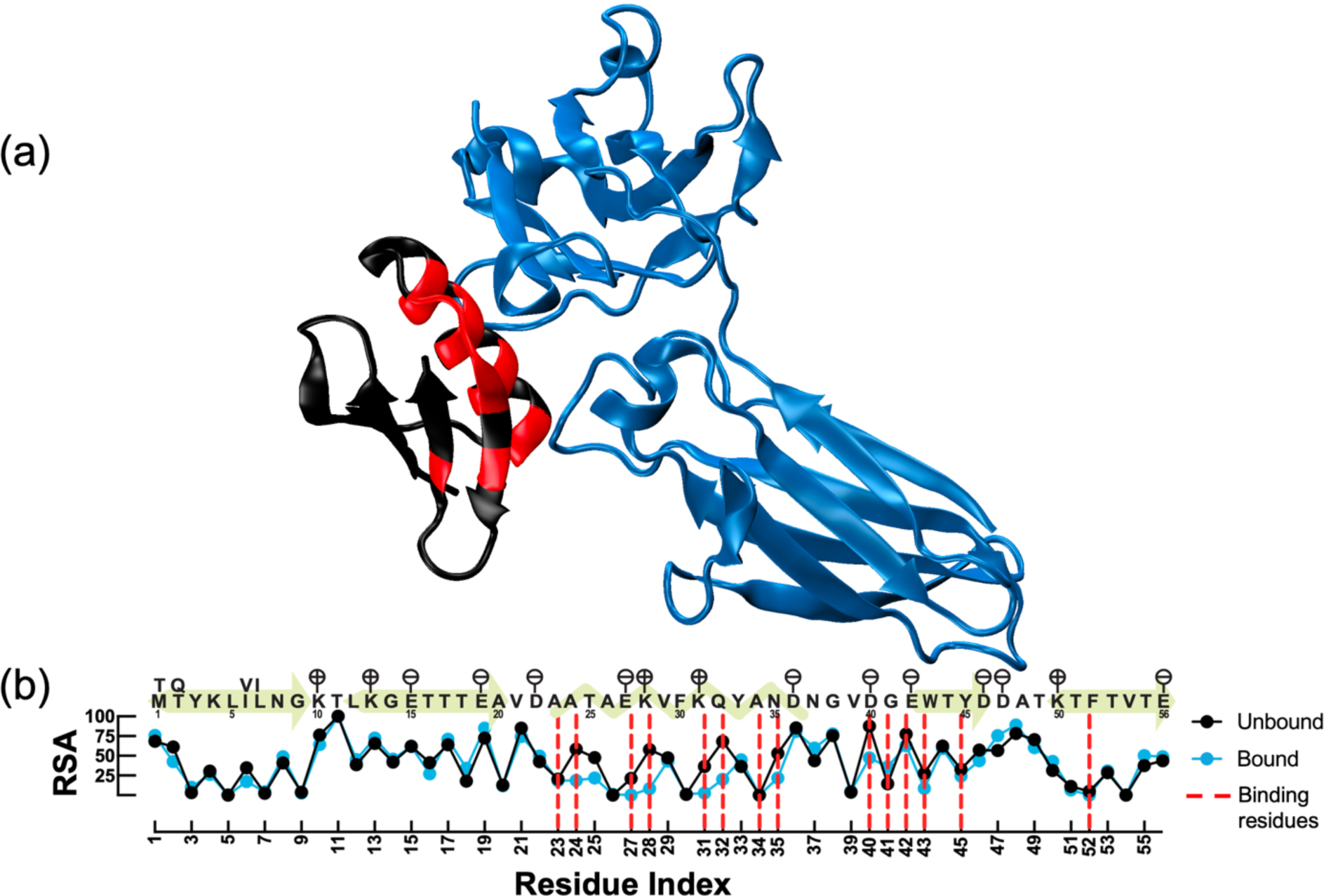
The C2 fragment of Streptococcal protein G (GB1, in black/red) partnered with the FC domain of human IgG (IgG-Fc, in blue). **(a)** WT form of the complex is employed for visualization and analysis by using minimized 1FCC PDB-coded structure. GB1 and IgG-Fc have 56 and 206 residues, respectively. Binding region of GB1 (residues that have at least one heavy atom within 5 Å of IgG-Fc) is shown in red. **(b)** Sequence of GB1 is illustrated by signifying secondary structure (yellow patterns) and charged residues (flags above positions). The amino acid differences between the sequences of 1PGA (unbound GB1) and 1FCC (bound form) are noted for residue positions 1, 2, 6 and 7. Accompanying RSA is calculated for unbound/bound forms of GB1 after minimization in water. Residues at the binding region are indicated by red dashed lines. The solvent accessibility of residues 24, 25, 27, 28, 31, 32, 35, 40 and 43 decreases after the binding, while a slight increase occurs for residues 19, 41, 47 and 48. There are 9 hydrogen bonds between GB1 and its partner in minimized WT. Residues 27 and 28 have two hydrogen bonds per position, and residues 35, 39, 40, 42 and 43 have one each.

Aiming to fill this knowledge gap, we have scrutinized the conformations and the dynamics of GB1 for unbound and bound forms by using computational methods. In particular, we seek to understand the degree to which changing the side chain followed by minimization in the correct solvent environment describes the shifts in the average positions of all atoms of the protein. This approach finds root in previous work where we have shown that in proteins, local displacements in the three-dimensional structure that disturb hard degrees of freedom, such as bond lengths, are not restored to their initial coordinates upon minimization but are instantaneously perpetuated to the whole structure through the rearrangements of soft degrees of freedom, such as torsions and non-bonded contacts.^23^ In fact, this idea leads to an efficient conformational search method for small molecules.^24,25^ The so-called mutation-minimization (MuMi) algorithm based on this idea has previously been applied for alanine mutational scanning to successfully determine positions critical for the functioning of Hsp70.^26^

In this study, we employ GB1 (Figure 1) and construct all possible single mutants of GB1 by using the MuMi scheme,^26^ as well as AlphaFold2 predictions of the mutants followed by minimization (AFMi).^27,28^ We also perform an extension of 1 ns-long MD simulation (MuMi-Dyn) for 14 positions (266 mutations) that are in close proximity to the binding interface. To form a solid basis for comparison, we conduct 2-µs long MD simulations of unbound and bound WT forms to establish the conformational dynamics sampled by the system at equilibrium. To understand the effects of single mutations on binding, we compare MuMi, MuMi-Dyn and WT-MD simulations for unbound/bound forms. We find that the MuMi scheme generated structures carry enough information to explain the stability shifts due to point mutations. We also find that the hydrogen bond occupancies at the GB1 – IgG-Fc interface is the main fingerprint of fitness. Since MuMi is an efficient method that has the right level of detail to predict the conformational shifts in proteins due to point mutations, our findings pave the way for developing machine learning methods that can locate the conformations on the fitness landscape.

## METHODS

### Simulations

A summary of all simulations conducted in this work is presented in Table 1 and are briefly described here. We use 1PGA and 1FCC coded crystal structures from Protein Data Bank^29^ (PDB) as unbound and bound forms of GB1 protein, respectively; the complex is displayed in Figure 1a and its sequence with the charged residues and mutated sequence positions are displayed in Figure 1b, top row. In the MuMi scheme, to construct 1064 single mutations (56 positions × 19 amino acid substitutions), we employ ProDy^15^ and Visual Molecular Dynamics (VMD).^30,31^ We then solvate the proteins in a cubic water-box that has a minimum distance of 10 Å to the protein in every direction so that a protein atom and another on its image are separated by at least 20 Å between. We add KCl salt to maintain the isotonic (0.15 M) concentration. Minimization of 10,000 steps is performed by using Nanoscale Molecular Dynamics (NAMD2) software, with CHARMM36 force-field.^32,33^ Alternatively, the prediction of protein structures belonging to single mutations is done by using ColabFold that uses AF as a base with a different MSA algorithm.^27,28,34^ We construct one model for each single mutation and use the same minimization procedure as in MuMi for the relaxation of structures in water; hence, this pipeline is named as AlphaFold2 and Minimization (AFMi). We also perform two duplicate 1-µs long MD simulations for unbound and bound forms of WT GB1 at 310 K constant temperature and 1 atm constant pressure on the minimized WT structures. We again use NAMD2 and CHARMM36 force-field for the MD simulations.^32,33^ Furthermore, 14 amino acid positions (14 x 19 = 266 mutant structures) are selected from the residues that have high probability to be located at the binding region according to the MuMi results (see Results and Discussion). 1-ns-long MD simulation extensions for these mutant forms are conducted by using the same pipeline as MD simulations for the WT. However, due to excess energy occurrence after the insertion of mutant side chains, we are unable to perform MD simulations for D40L, E42H, E42M, E42F and E42Y single mutant structures. By omitting these mutations, we produce 261 systems by using MuMi-Dyn method for further analyses.

**Table 1.** List of simulations conducted in this work.

| Abbreviation | Methodology | Number of mutations | Length of each MD | Total length of simulation |
| --- | --- | --- | --- | --- |
| WT unbound MD | Classical MD | - | 1 μs (two replicates) | 2 μs |
| WT bound MD | Classical MD | - | 1 μs (two replicates) | 2 μs |
| MuMi unbound | Mutation + minimization | 1064 | - | - |
| MuMi bound | Mutation + minimization | 1064 | - | - |
| AFMi unbound | AF prediction + minimization | 1064 | - | - |
| MuMi-Dyn unbound | MuMi + 1ns MD relaxation | 261 | 1 ns | 261 ns |
| MuMi-Dyn bound | MuMi + 1ns MD relaxation | 261 | 1 ns | 261 ns |

### Analyses of structures

Solvent accessible surface area (SASA) and hydrogen bonds are computed by VMD plugins.^30^ The hydrogen bond definition used is 3.0 Å donor-acceptor distance and 20° for the hydrogen bond angle. FreeSASA^35^ software is used for relative solvent accessibility (RSA) analysis to investigate the changes in solvent interaction profiles for each residue belonging to a protein structure. To calculate probability of being located at the binding region, we compute distances of all heavy (non-hydrogen) atoms between GB1 and its partner; then, if any two atoms of these two compounds are closer than 5 Å,^12^ the residue index on GB1 is saved as one occurrence of being located at binding region. All occurrences are calculated for WT structure, MuMi, MuMi-Dyn and WT-MD simulations and displayed as probability density functions. Residues described as allosteric for GB1 (residues 9, 12, 14, 30, 33, 37, 39 and 56) have been taken from a recent study.^12^

## RESULTS

### GB1 – IgG-Fc interface is maintained by dynamical hydrogen bonds

The interface was defined by a RSA analysis which was previously used to categorize the surface and core residues of GB1.^12,36^ We calculate the RSA of each amino acid from the minimized WT structures and find that the RSA values and the ensuing definition of binding residues differ from the previous work where only the crystal structure was used for a similar analysis (Figure 1).^12^ This is because the crystal structures might represent artificial lattice restraints which are released upon minimization in the solvent environment. Residues near the binding region have propensity to interact less with water (Figure 1b), in particular, the solvent accessibility of residues 24, 27, 28, 31, 32, 35, 40 and 43 decreases in the bound form.

We next study the dynamics of unbound and IgG-Fc bound GB1 through duplicate MD simulations, each of 1 *µ*s length. The RMSD and RMSF plots for these runs are presented in Figure S1. Both x-ray structures represent equilibrium states with less than 2 Å RMSD from the initial state of the system throughout the 2 x 1 *µ*s length of the simulations. Increased RMSF values in the bound state for the residues distal to the binding interface reflects a general property of proteins.^37–39^ While this increased flexibility is usually accompanied by the rigidification of the binding site residues, in GB1 we find the latter effect to be minimal, because the interface is made up of secondary structural elements (Figure 1b).

A closer look at the interactions at the interface of GB1 upon binding lends further clues on the dynamics. According to the mutational scanning experiments of Olson *et al*.,^16^ the least mutatable positions are E27, K31, and W43. All three of these residues are classified as ‘interface’ residues due to the definition that any of their heavy atoms is within 5 Å of the binding partner.^12^ When effects of binding stability and folding stability are differentiated in these data, it was found that the stability of these residues all stem from the former.^9^ Of these, D27 and K31 are positioned at the binding interface at a distance which would enable them to form a salt bridge. However, our MD simulations reveal that the intramolecular D27-K31 is a loose salt bridge in the *apo* form while it is completely stabilized upon binding (Figure 2a). This stabilization is caused by the strong K28-E380 and K28-E382 interactions at the interface (Figure 2b,c). Meanwhile, W43 performs a dual function. On the one side, with its bulky side chain, it partially occupies the hydrophobic core of the protein, being in contact with L5, A34, F30, F52 and V54. On the other side, it interfaces with the hydrophobic carbon tail of the K31 side chain close to the backbone, placing its NH_3_^+^ group in the correct orientation for the E27-K31 interaction mentioned previously. Thus, these residues are excellently positioned to stabilize the interface via the K28 interactions.

**Figure 2.**
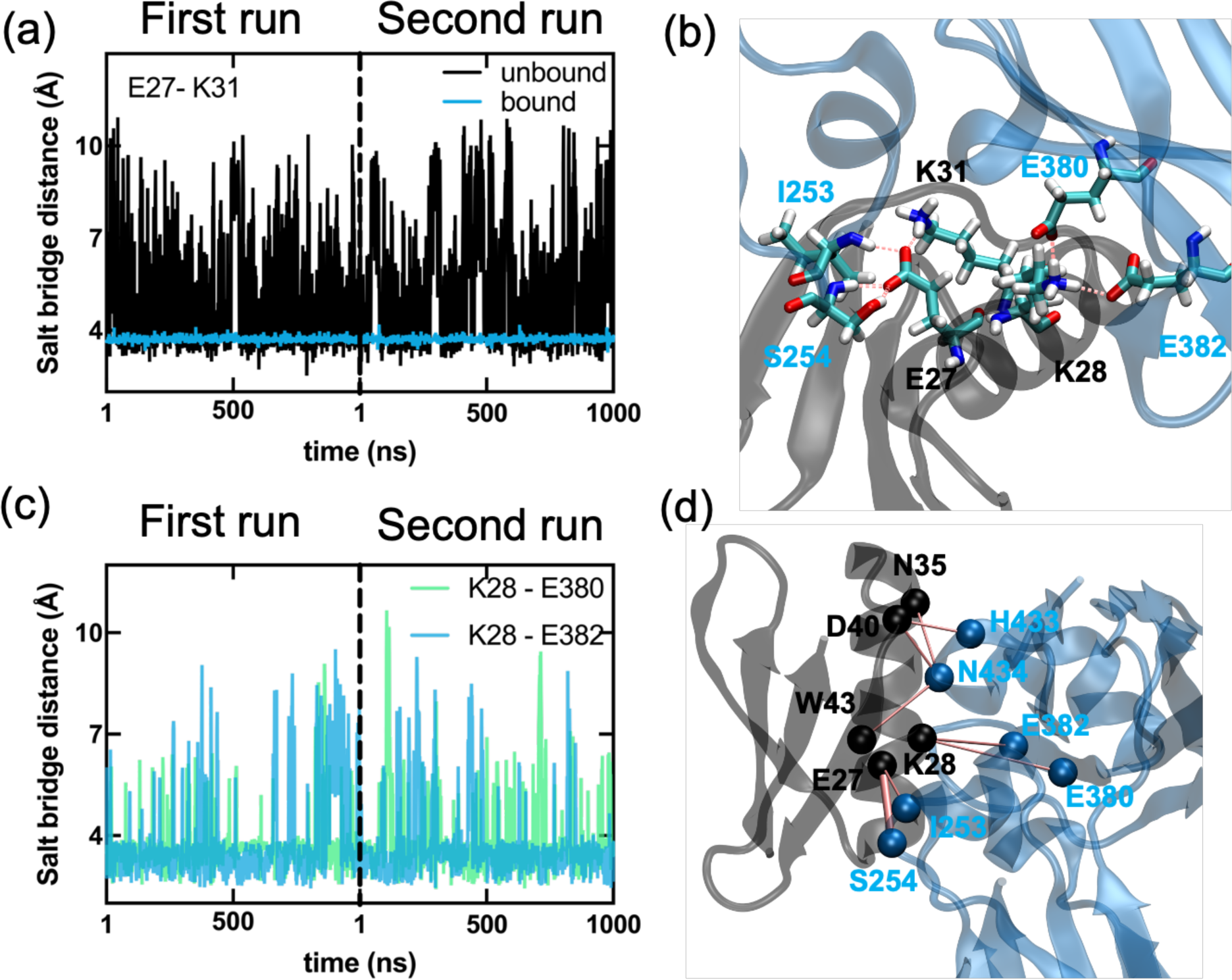
Key distances from WT MD simulations; two independent 1µs runs for each of the unbound and bound forms of GB1. **(a)** E27-K31 distance in the apo structure is highly variable (black) but is completely stabilized upon binding (blue). **(b)** The coordination of salt bridges occurs between residues E27 and K28 of GB1 and residues I253, S254, E380 and E382 of IgG-Fc. Here, K31 stabilizes E27 with a salt bridge, leading to successful binding. The annotated residues are shown in licorice, and the salt bridges are depicted as dashed pink lines. **(c)** Stabilization at the interface is due to dynamical interactions of K28 from GB1 with E380 and E382 from IgG-Fc. **(d)** The network of interactions at the GB1 and IgG-Fc (see also Table 2).

**Table 2.**
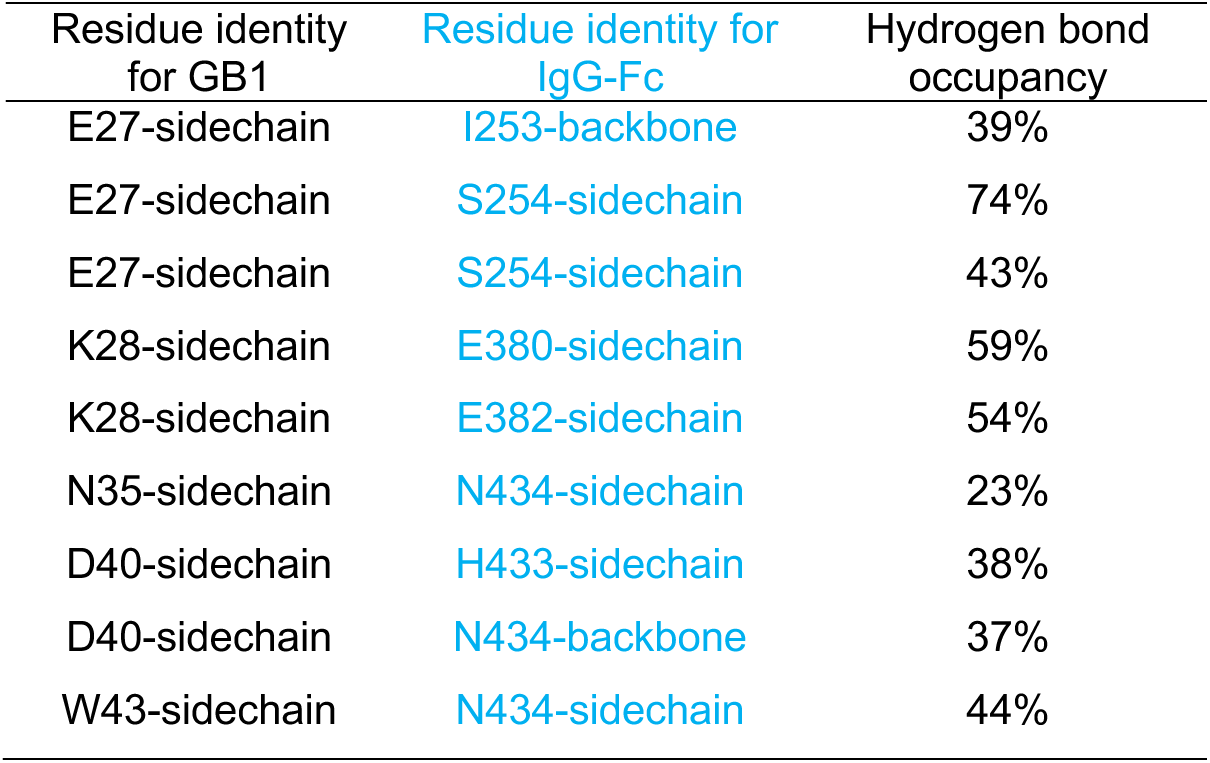
Hydrogen bond occupancies from 2 µs MD simulations at the GB1/IgG-Fc interface.

In Table 2, we list the 9 hydrogen bonds between GB1 and IgG-Fc with occupancies greater than 20 % in the total of 2 µs WT-MD runs. We plot the dominant hydrogen bond network that achieves specific binding in Figure 2d.

### MuMi-generated structures encapsulate the key aspects of binding fitness information

We next seek if mere mutation of positions followed by minimization produces plausible structures that represent conformational shifts accompanying the side chain replacements. In Figure 3, we display three example mutations demonstrating that while the change introduced is local, long-range shifts in positions are achieved due to rearrangements in the soft degrees of freedom, which may lead to RMSD values exceeding 1 Å.^24,25,37^ In fact, we find that the plain count of the number of hydrogen bonds maintained at the interface following the MuMi scheme, where we display the scanning of the 56 positions with 19 amino acid substitutions in 1064 positions in Figure 4a, is a very good predictor of fitness for this system (Figure 4b).

**Figure 3.**
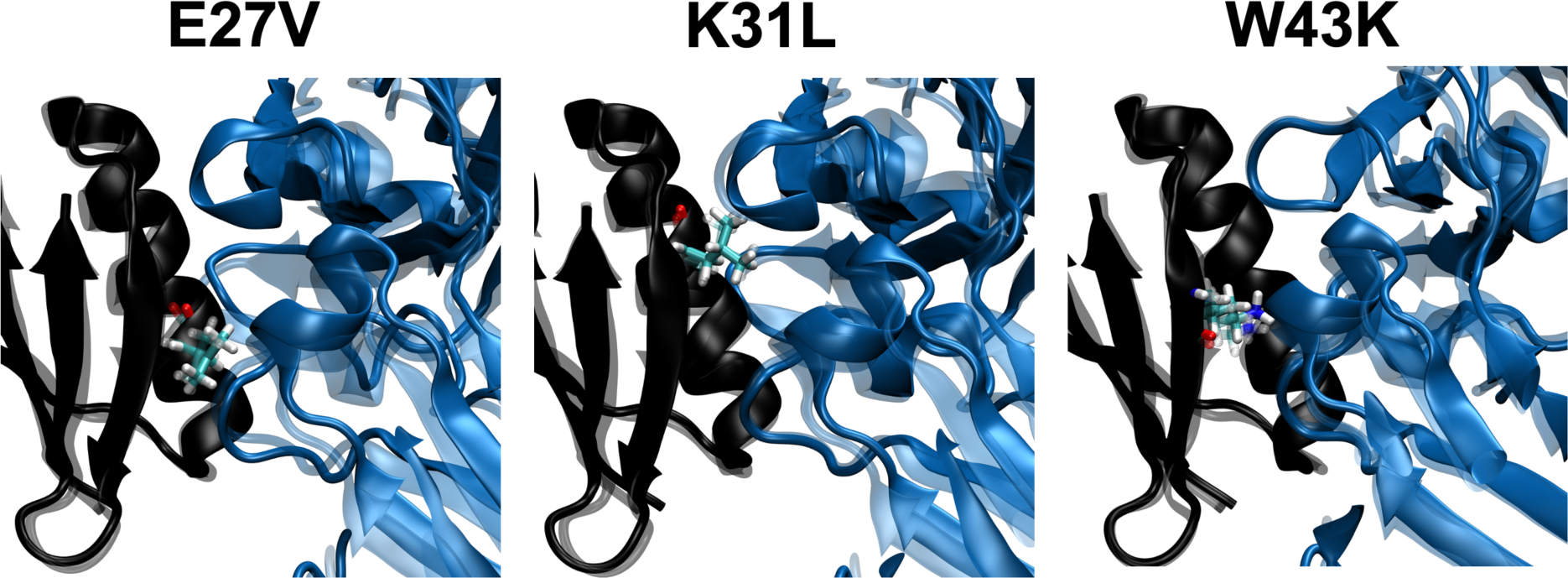
The effect minimization for E27V, K31L and W43K mutants. GB1 and IgG-Fc are shown in black and blue. Mutated residues are shown as licorice with atom coloring. Initial (mutated WT crystal structure, 1fcc PDB-coded) structure is illustrated as transparent; on the other hand, final minimized structures are illustrated in opaque colors. Heavy atom RMSD between the initial and final structures is 1.15, 1.07 and 1.11 Å for the respective MuMi replacements.

**Figure 4.**
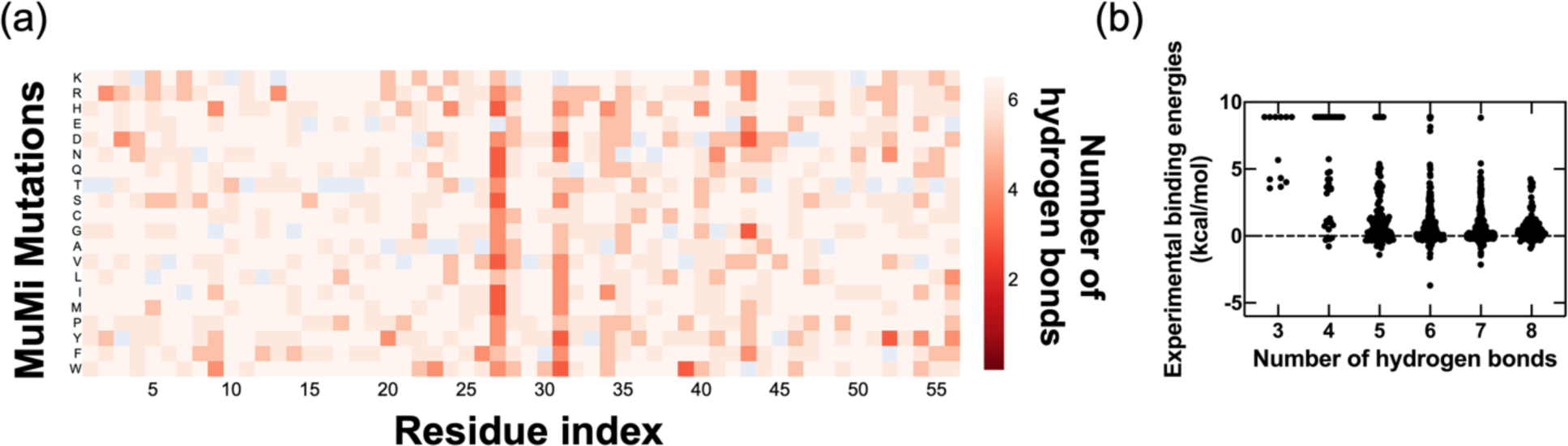
**(a)** Heatmap of number of WT intermolecular dominant hydrogen bonds for all MuMi-produced conformations. **(b)** The number of WT dominant is compared with the experimental binding energies are shown on the right. The experimental binding energies are capped at 9 kcal/mol and any mutation exceeding this amount of destabilization is reported through this value.^9^

We note the correspondence between the low stability caused by mutations at positions 27, 31 and 43 discussed in the previous subsection to be well described by the loss of the interface interactions. Interestingly, the K28 residue itself which establishes two stable hydrogen bonds in the WT (Figure 2b,c) is relatively more tolerant to single residue replacements which exactly follows the experimental findings,^12^ pointing to an intricate balance of interactions at the interface.

### Conformational variability in mutant protein structures generated by MuMi vs. AFMi

Since generation of point mutation structures may also be obtained by alternative approaches, we next compare our MuMi and AF predicted structures. We performed MuMi and AFMi calculations for all single mutants and calculated the RSA for final structures. Following recent work where classification of residues based on solvent accessibility without any measurement of binding energies showed promise,^12,13,15^ we use RSA as a measure of the effect of mutations on the amino acid positions. We find that not only the means, but the standard deviations (*σ*) and probability density functions (PDFs) are informative about the conformational effects of mutational perturbations. The MuMi and AFMi predicted structures are on average very similar in terms of ⟨RSA⟩ (Figure 5a). However, comparison of their variance shows that there are significant differences in the predictions for positions 39, 41, 46 and 56 (Figure 5b-c). Residue 41 is known for its position at the binding interface (Figure 1b) without taking part in the hydrogen bond network of Table 2. Residues 39 and 56 were reported as allosteric.^12^ The RSA probability distributions for these residues are displayed in Figure 5d (see Figure S2 for all positions). Positions 39 and 56 sample a wider range solvent exposure in AFMi predictions. Conversely, residue 46 has two states of different solvent exposure in MuMi structures, while it occupies only one state in those from AFMi. Position 41 samples a variety of states in both approaches, but more so in MuMi than AFMi; since this is a glycine in the WT and is located on a loop, perturbations cause significant impact on its conformations. Finally, we find that the total SASA distribution of AFMi generated structures is shifted into larger values than those from MuMi (Figure 5e); i.e. AF predicts slightly expanded structures for GB1.

**Figure 5.**
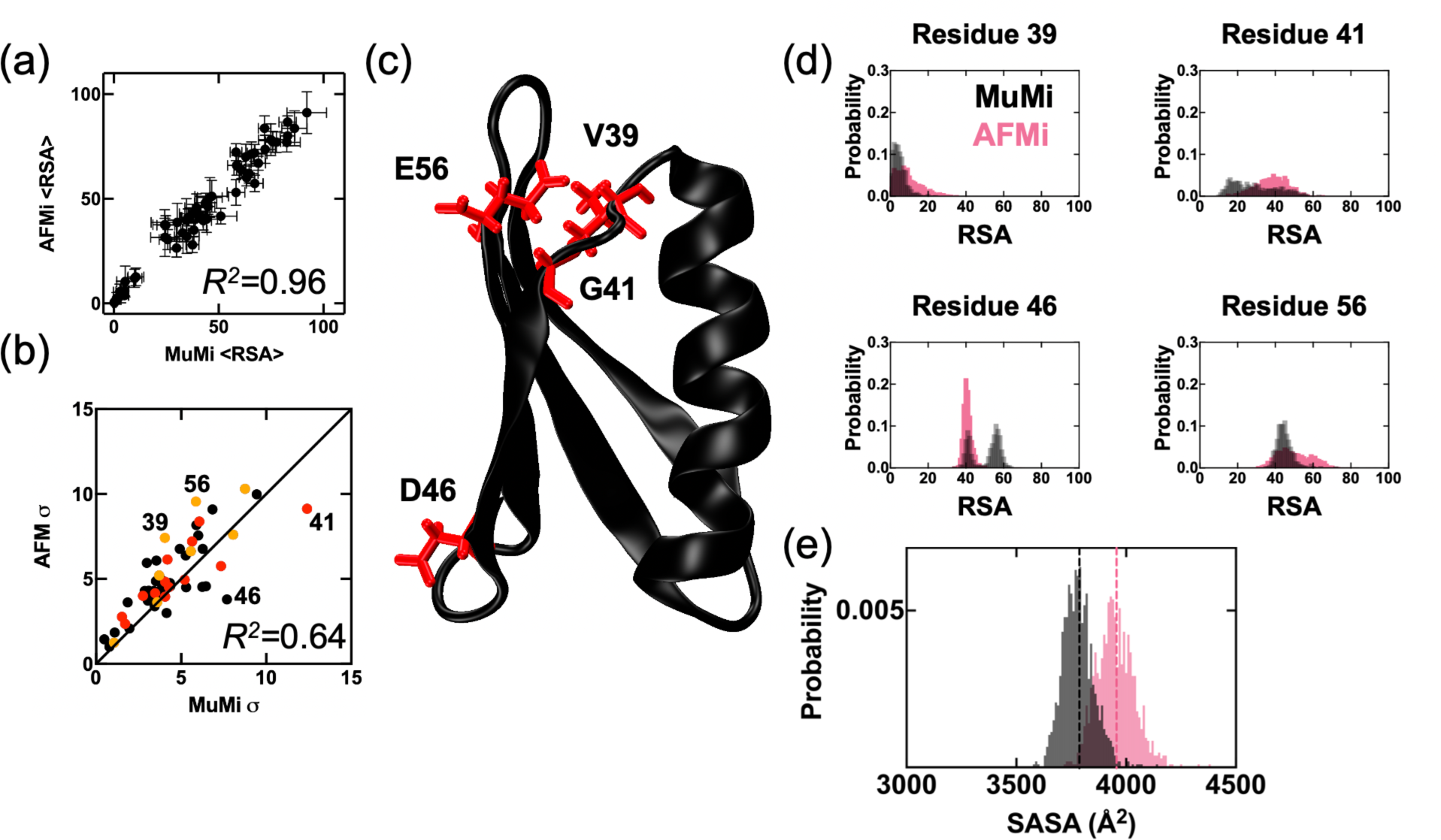
**(a)** RSA results of all-possible single mutation conformations (56 positions × 19 amino acid substitutions) generated by MuMi and AFMi schemes for unbound GB1. ⟨RSA⟩ shows the average effect of mutational perturbations to that amino acid position through the single mutations. ⟨RSA⟩ results of single mutations for MuMi and AFMi are also very similar, which indicates that mutational insertions to the crystal structure and AF-predicted mutant conformations are very similar on average. **(b)** Comparison of standard deviation (*σ*) of RSA for MuMi and AFMi showing the importance of probability distributions over averaging to scrutinize the effects of single mutations. Binding region residues from the minimized WT complex are shown in red, and allosteric residues from a previous study^12^ are shown in orange. **(c)** Outliers from (b) are visualized on the WT unbound-GB1 structure. **(d)** Probability density functions (PDFs) of RSA values of single mutations of outlier positions. (e) Total SASA of the proteins from AFMi (pink) and MuMi (gray).

In general, the difference between MuMi and AFMi indicates that, to construct conformations representing single mutations, starting from the crystal WT structure is a sound method to understand the mutational impact. In contrast, the conformational differences obtained via AFMi emerge from the varying MSAs (Figure S3) and training the weights of AF; they are therefore likely carry additional sources of variations.^40^

### MD simulations and mutational scanning provide consistent insights into the functional dynamics of the protein at the molecular level

The relationship between conformational variability observed in µs-long MD simulations and that derived from mutational scanning studies is an aspect of interest in protein dynamics and function analysis. Thus, we analyze the overlap between the range of conformations sampled by the WT structure and those covered via single residue replacements. We analyze the 2-µs-long MD simulations both for the unbound and the bound forms of GB1 and compare our findings with MuMi by focusing on three factors: **(i)** The number of hydrogen bonds between GB1 and its partner; **(ii)** RSA differences of unbound/bound forms of GB1; and **(iii)** the probability of finding a residue at the binding region. The hydrogen bonds inform on the robust interactions between GB1 and IgG-Fc, and by interpreting the solvent accessibility we understand the water mediated changes.^14,41^ Additionally, the assessment of GB1 residues located in the binding region is important to investigate the interactions between the protein and its binding partner.^12^ We find that for all single mutations, 14 positions have a higher probability of being located at the binding region than the WT (Figure 6a). We apply 1-ns-long MD simulations following MuMi for this subset, culminating 261 MuMi-Dyn simulations and confirm that the hydrogen bond distribution of 1-ns-long extension does not significantly differ from that of the WT-MD simulations.

**Figure 6.**
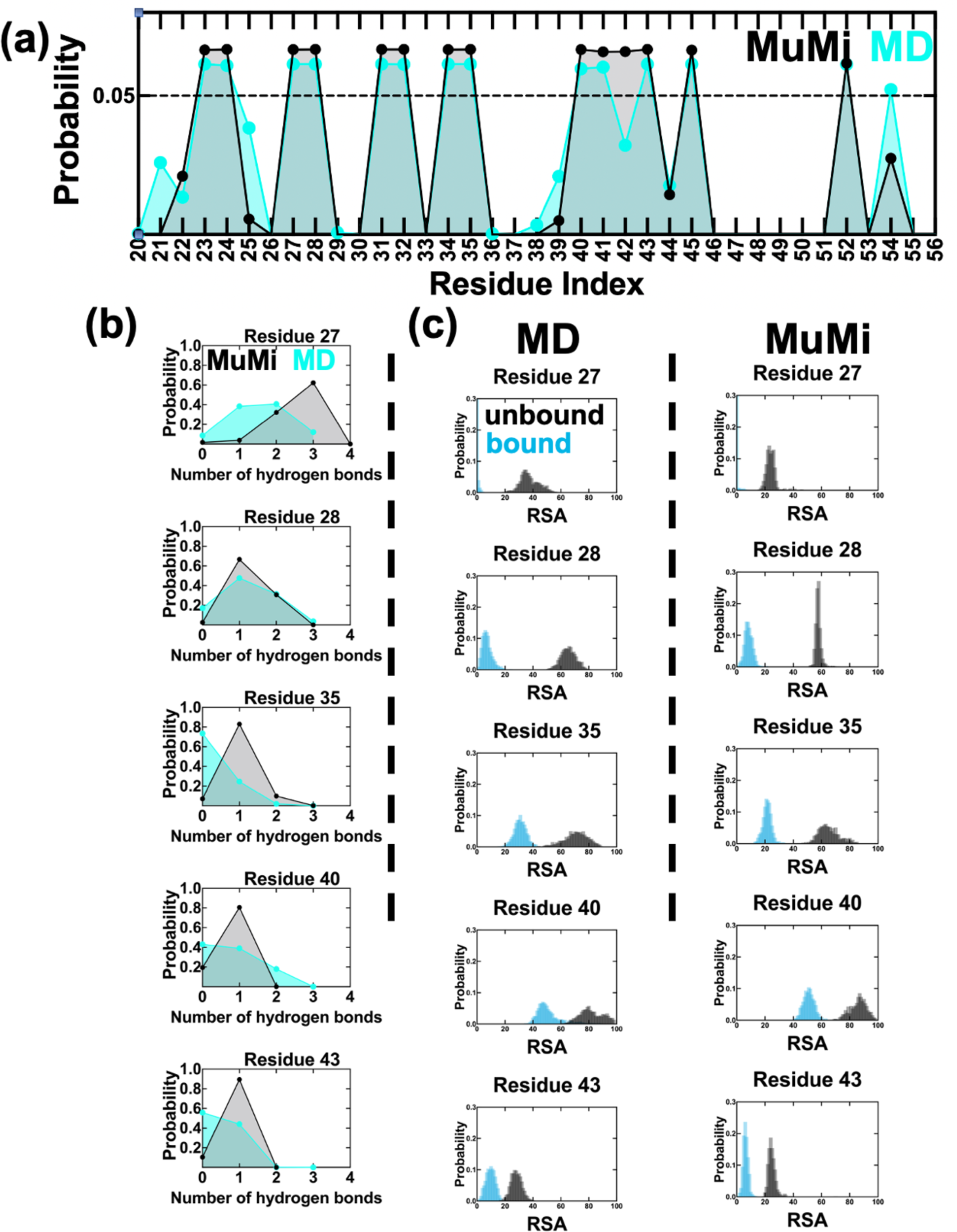
Assessment of GB1 binding by using MuMi and MD simulations. **(a)** Probability of residues being located at the binding region for mutations (MuMi) and MD simulations. Residues 1–19 are not shown due to insignificant probabilities. In MD results, residues 21, 25, 39 and 54 have higher propensity to stay at the binding region compared to MuMi, while residue 42 has lower probability. **(b)** PDFs of hydrogens bonds established between GB1 and IgG-Fc for residues listed in Table 2, calculated for MuMi and MD simulations. Residues tend to have a lower number of hydrogen bonds in MD simulations in contrast to MuMi. **(c)** PDFs of RSA are calculated for unbound/bound forms for the same residue set (see Figures S6, S7 for PDFs of the complete set of residues of unbound/bound for MuMi and WT-MD).

In Figure 6b, we compare the probability distribution of the hydrogen bonds whose occupancies are listed in Table 2 with those obtained from MuMi (a comprehensive picture for all residues that display at least one hydrogen bond is in Figure S4). As a result of thermal fluctuations in the MD simulations, the total number of hydrogen bonds found at the interface averaged over the duration of the WT-MD is less than those found in the ensemble of minimized structures of MuMi (Figure S5a,b). Adding relaxations via 1 ns long short dynamics on the mutants of the 14 residues located near the binding site releases some of the hydrogen bonds in the minimized structures and covers a range similar to that of MD-WT (Figure S5a,c).

As a result of the less tight binding at the interface, the protein SASA of unbound/bound conformations for WT-MD are higher than MuMi (Figure S6a,b). While this reflects into the RSA of individual residues (see examples in Figure 6b) MuMi predicts the local effects remarkably well and the coverage of single residue replacements is similar to that of the WT ensemble collected under dynamical conditions whereby large-scale motions and the impact of water are pronounced by the MD simulations. PDFs of all hydrogen bonds and RSA analyses for MuMi and WT-MD simulations are displayed in Figures S7 and S8.

We find that SASA and total hydrogen bonds in MuMi-Dyn are similar to those from WT-MD, since the more dominant water interactions in MD simulations modulate these factors (Figures S5, S6). Residues also generally have a lower number of hydrogen bonds as in WT-MD simulations in MuMi-Dyn method (Figures S5c, S9); however, residues 40 and 41 have lower probability of being located at the binding region compared to MuMi and WT-MD (Figure 6a vs. Figure S10). Considering that MuMi-Dyn was applied only on binding region residues, it is interesting to have a similar regime with the WT-MD simulations with these biased mutational perturbations. To compare water interaction of residues for MuMi, WT-MD and MuMi-Dyn, we compare ⟨RSA⟩ and *σ* of RSA of each method for unbound/bound GB1.

### Similarity of water interaction regime for MD simulation of WT and the single mutations

Considering mutational insertions and their physical features, such as hydrophobicity, volume and charge, even with the drastic differences between the amino-acid type of WT and the mutation, the effects of single mutations may fall within stability margin.^36^ Therefore, we observe slight differences between the conformational space of WT and the single mutants through the MD trajectories and expect an overall similarity.

To this end, we compare RSA values of unbound/bound GB1 for MuMi, WT-MD and MuMi-Dyn, and find that the methods with MD simulations have wider RSA PDFs (Figure S7, S8, S11). Overall similarity between ⟨RSA⟩ of MuMi, WT-MD and MuMi-Dyn indicates that water accessibility of residues is conserved through the single mutations (Figure 7). Note also that *σ* of RSA values are significantly similar for WT-MD and MuMi-Dyn results. (Figure 7 e, f). Like the comparison of MuMi and AFMi, analyses of unbound and bound forms of GB1 do not differentiate the allosteric and binding residues. This result indicates that GB1 achieves the function by modulating slight changes in the whole system, which was also observed in PDZ3.^41,42^

**Figure 7.**
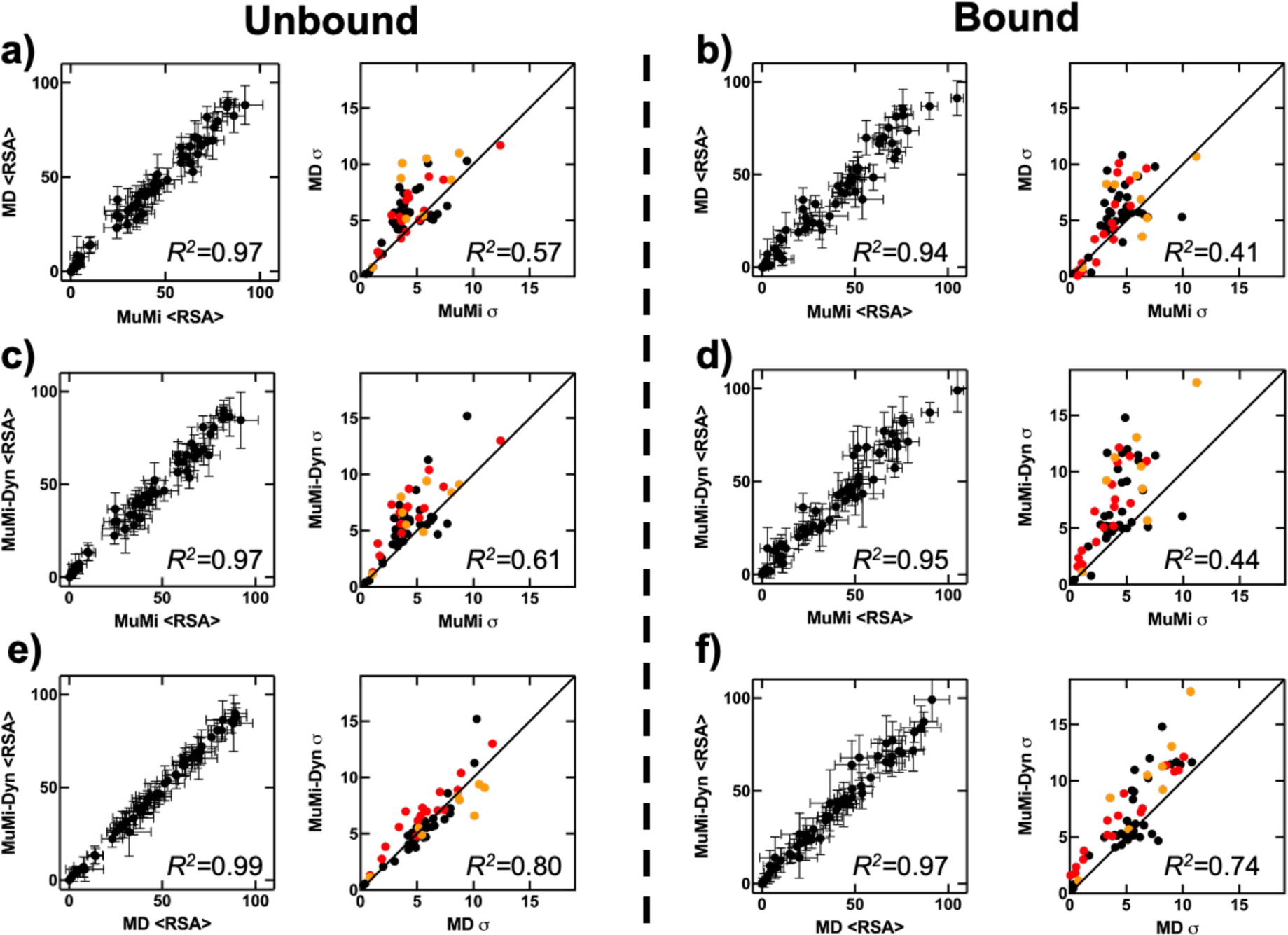
Comparison of ⟨RSA⟩ of unbound/bound form for MuMi, MuMi-Dyn and WT MD simulations. Allosteric residues and binding residues are indicated with orange and red, and *R*^2^ values are written on graphs. a, b) The ⟨RSA⟩ and σ values of MuMi and MD simulations for unbound/bound. c, d) The ⟨RSA⟩ and σ values of MuMi and MuMi-Dyn for unbound/bound. e, f) The ⟨RSA⟩ and σ values of WT-MD and MuMi-Dyn simulations for unbound/bound. (see Figure S4, S5 and S9 for PDFs of all residues by MuMi, MuMi-Dyn and WT-MD simulations).

In the minimization process of MuMi, the effect of the local mutations is propagated to the instantaneous interactions. Conversely, the thermal fluctuations and solvent interactions are sampled via MD simulations leading to conformations reflecting the mutational characteristics more significantly. Interestingly, the biased mutations in the binding region and the coverage from WT-MD simulations are very similar in terms of ⟨RSA⟩ (Figure 7 e, f). These results indicate that, although, there are exceptions like ‘deleterious’ mutations, most of the single mutations modulate only the conformational space of protein structures. By fine-tuning the conformation space in varying fractions of time, the impact of single mutations on protein function/binding emerges. Thus, with larger sample sizes and longer MD simulations, it might be possible to investigate the conformational space of possible single mutations, without physically applying amino acid mutation on a protein structure.

## DISCUSSION

DMS provides valuable insights into the structure-function relationships of proteins, the molecular basis of disease, and the mechanisms of protein evolution. It is widely used in protein engineering, drug discovery, and systems biology research to understand and manipulate protein function with high precision. Computational prediction of the effect of point mutations on the conformational landscape of proteins has the potential to overcome a crucial bottleneck in developing an understanding of the point mutational landscape of proteins.

To construct the single mutations of GB1, we compare MuMi and AFMi; the latter is an MSA-based structure prediction method,^28^ and the former is a crystal structure-based method.^26^ Overall, this comparison suggests that using the crystal WT structure as a starting point for modeling single mutations is a reliable method for understanding their impact. Conversely, the conformational variations observed with AFMi are due to differences in the MSAs and the tuning of AlphaFold’s weights, which may introduce additional sources of multiplicity. Hence, we find that, it is more prudent to perform mutagenesis by using a WT crystal structure.

The most striking finding of this work is that MuMi, a computationally feasible approach to generating computational DMS structures, provides enough information to reproduce the most important aspect contributing to the fitness of GB1; i.e. binding energy (Figure 4). While we find a single feature defined as the number of hydrogen bonds at the interface already correlates well with the fitness, this observation paves the way to using MuMi generated structures as the input to machine learning algorithms to predict fitness landscapes through computational DMS.

The binding of GB1 is further scrutinized by focusing on three factors, that are hydrogen bonding of GB1 to IgG-Fc, probability of being located at the binding region and RSA analysis. Our results comply with a previous study,^17^ where residues 39, 40 and 41 were experimentally shown to have important effects on binding of GB1. The effects of single mutants are mostly subtle, and the conformation distributions do not discriminate between allosteric and binding residues. By employing these three analyses, we describe the most important elements of binding.

## ASSOCIATED CONTENT

### Data availability

The code used for MuMi scheme can be found at https://github.com/midstlab/MuMi_scheme. All raw data and analyses codes are deposited at https://zenodo.org/records/11399775.

### Supporting Information

Figure S1. The RMSD and RMSF results of MD simulations for unbound/bound GB1; Figure S2. RSA probability distributions from MuMi and AFMi; Figure S3. MSA depth for predicted AF structures; Figure S4. Probability of hydrogen bonds between GB1 and the partner for MuMi and WT MD simulations; Figure S5. Probability distribution of total hydrogen bonds between GB1 and IgG-Fc; Figure S6. Probability distribution of SASA for unbound/bound GB1; Figure S7. Probability of RSA of unbound/bound GB1 for MuMi; Figure S8. Probability of RSA of unbound/bound GB1 for WT MD simulations; Figure S9. Probability of hydrogen bonds between GB1 and the partner for MuMi-Dyn; Figure S10. Probability of being located at the binding interface for MuMi-Dyn.

## AUTHOR INFORMATION

### Corresponding Author

**Tandac Furkan Guclu** - Faculty of Engineering and Natural Sciences, Sabanci University, 34956, Istanbul, Turkey; ORCID: 0000-0002-2516-1922;

### Authors

**Ali Rana Atilgan** - Faculty of Engineering and Natural Sciences, Sabanci University, 34956, Istanbul, Turkey; ORCID: 0000-0003-0604-6301

**Canan Atilgan** − Faculty of Engineering and Natural Sciences, Sabanci University, 34956, Istanbul, Turkey; ORCID: 0000-0003-0557-6044

### Author Contributions

T.F.G. designed the research, conducted the computer experiments, analyzed results, interpreted data, wrote the paper, and constructed figures. A.R.A. and C.A. designed the research, guided the computer experiments and analyses, interpreted data, guided the structure and contents of the paper, and edited the paper. All authors have given approval to the final version of the manuscript.

## Acknowledgments

The numerical calculations reported in this paper were partially performed at TUBITAK ULAKBIM, High Performance and Grid Computing Center (TRUBA resources).

## TABLE OF CONTENTS GRAPHIC

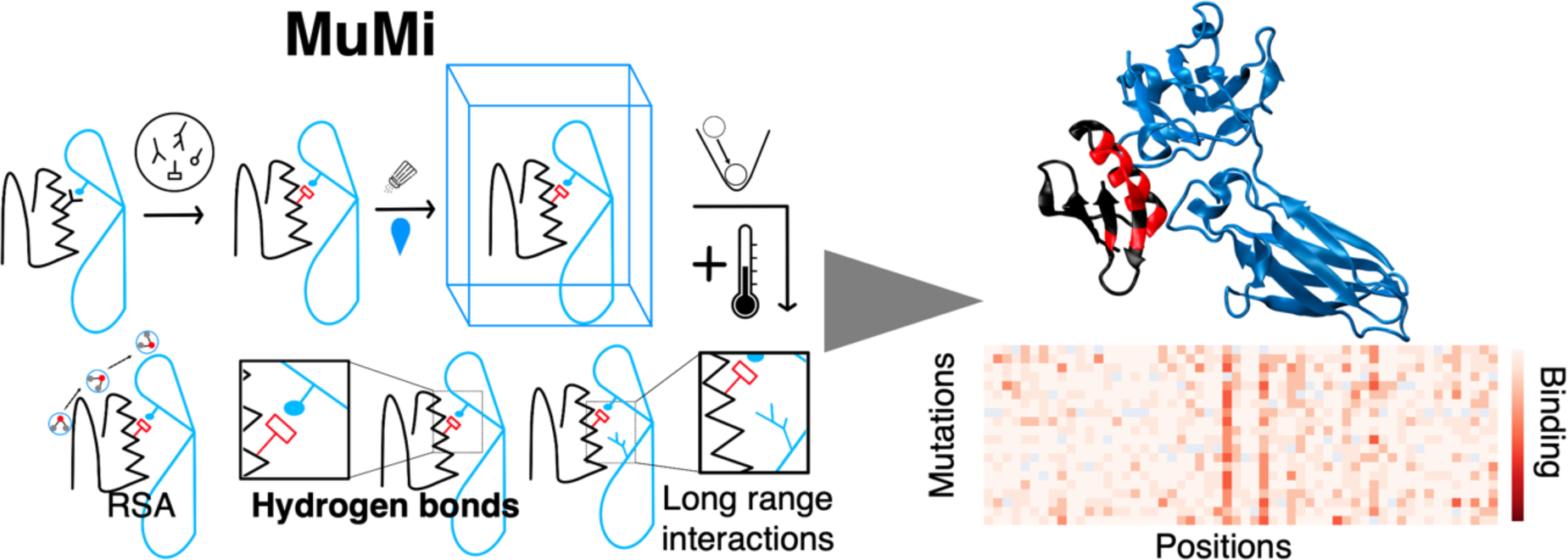

## Supporting Information

**Figure S1.**
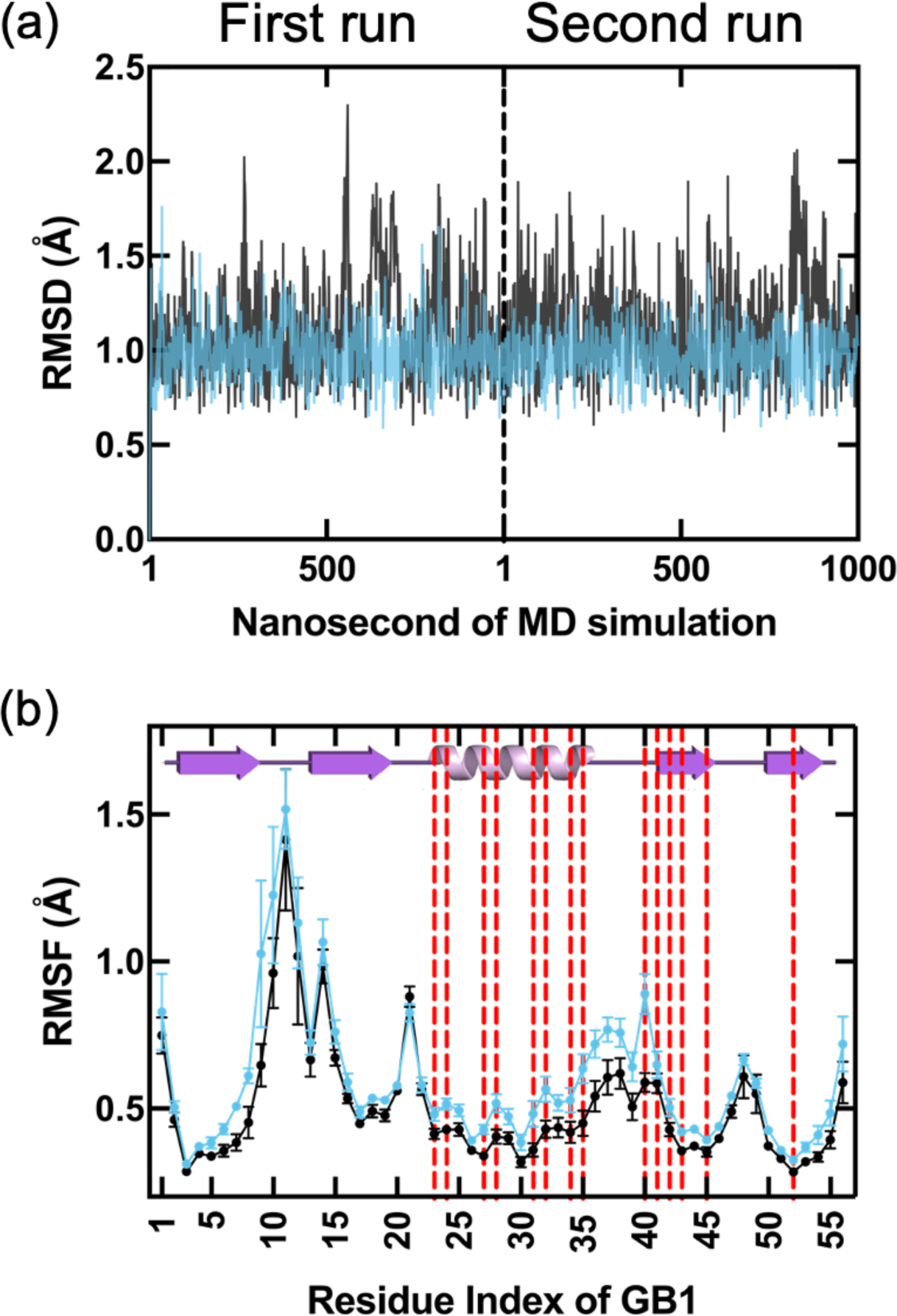
**(a)** Root mean squared deviations (RMSD) from the MD simulations of the unbound (black) and bound (blue) forms of the WT GB1. The two runs are appended with the dashed vertical line separating them. Reference structure is the respective minimized WT crystal structure. **(b)** Root mean squared fluctuations (RMSF) results for the same runs. Averages are taken over eight, 250 ns-long batches; mean and standard deviations are shown on graph. The binding residues of the minimized WT complex are indicated with dashed red lines.

**Figure S2.**
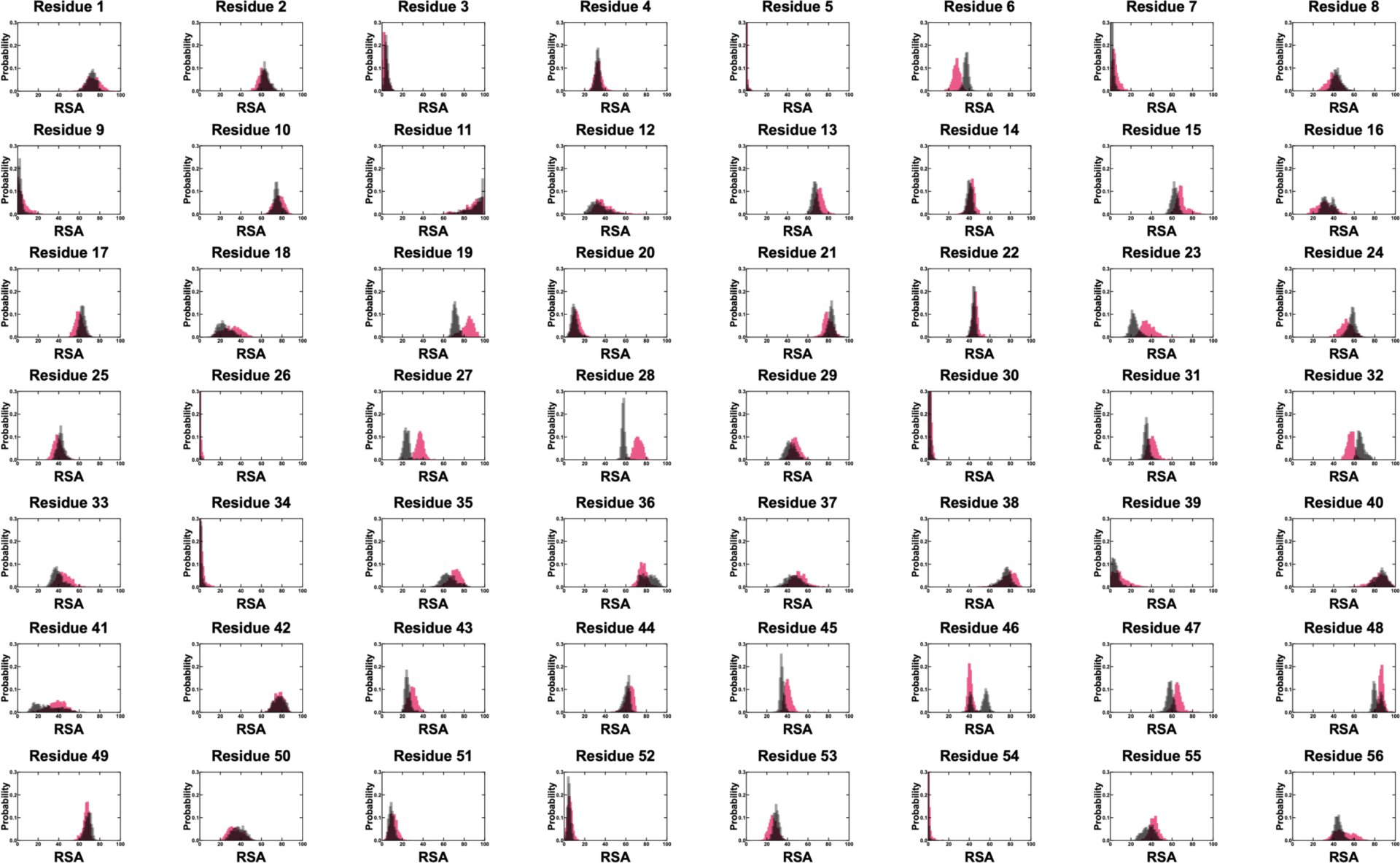
RSA probability distributions from MuMi (in black) and AFMi (in pink) for all 56 positions for unbound GB1.

**Figure S3.**
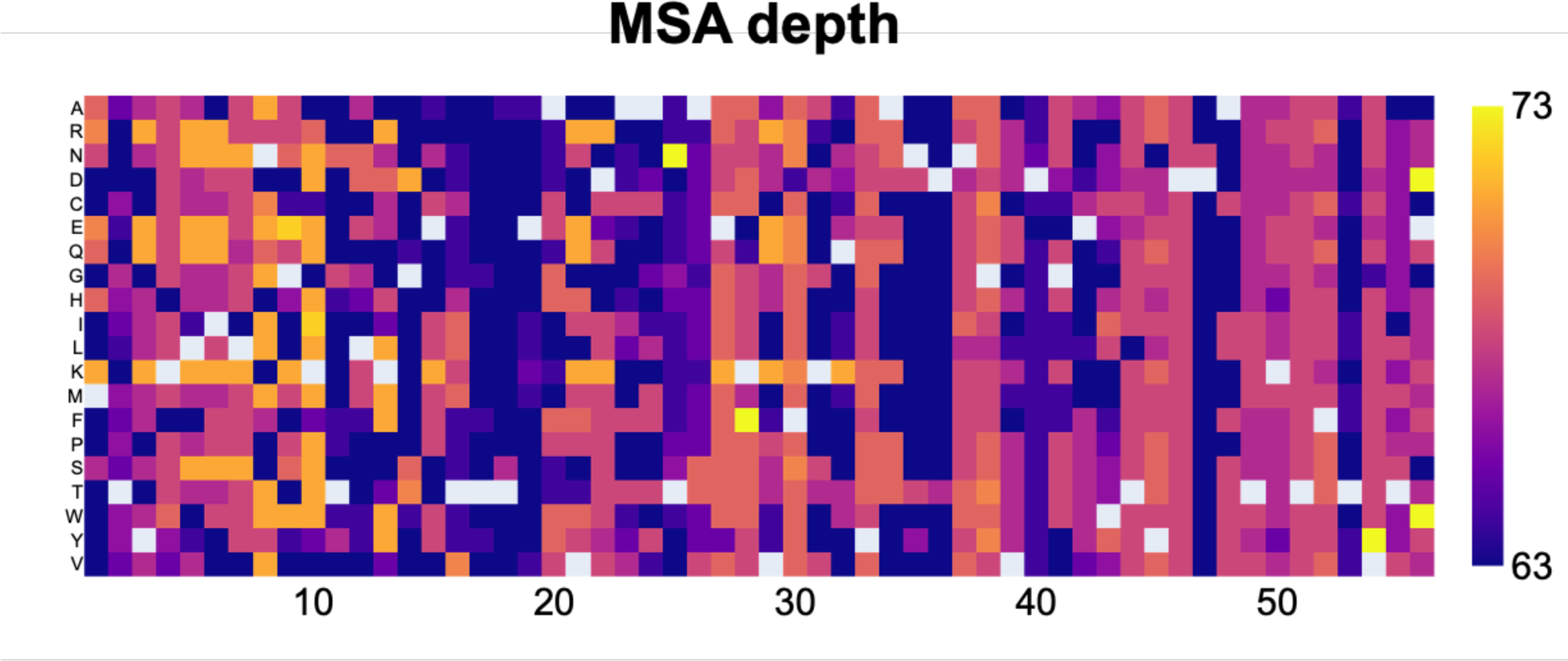
Multiple sequence alignment (MSA) depth for predicted AF structures. Dark blue and yellow correspond to 63 and 73, respectively. White squares indicate the self-mutations, which are not computed. MSA depth shows the number of unique protein sequences in an MSA that is used to predict structures. Difference between MSAs causes the variation between the predicted structures. MSA coverage has also been calculated, and the values change between 0.97–0.98.

**Figure S4.**
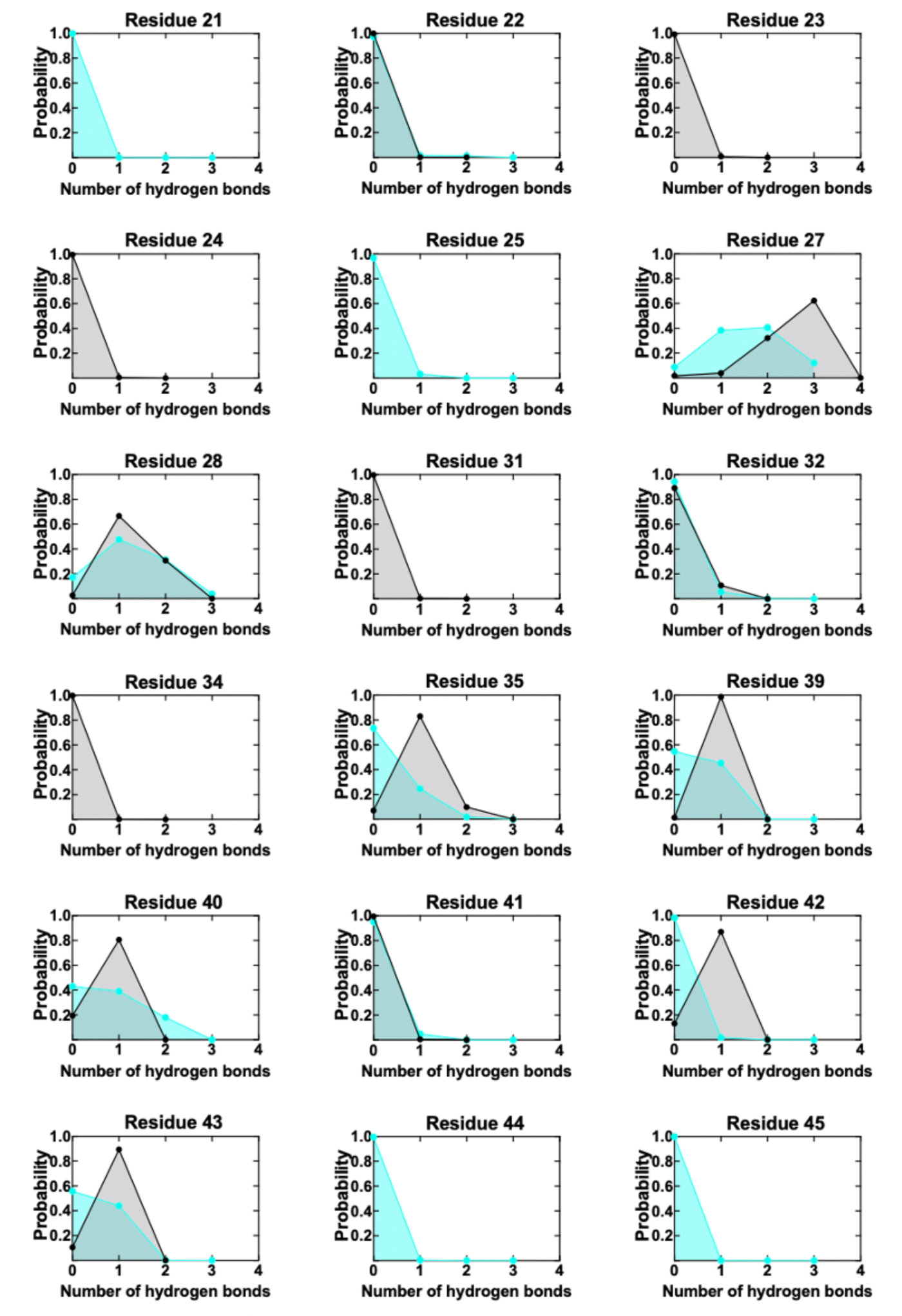
Probability of hydrogen bonds between GB1 and the partner for MuMi (in black) and WT MD simulations (in cyan). All residues having even a single occurrence of hydrogen bonds are shown.

**Figure S5.**
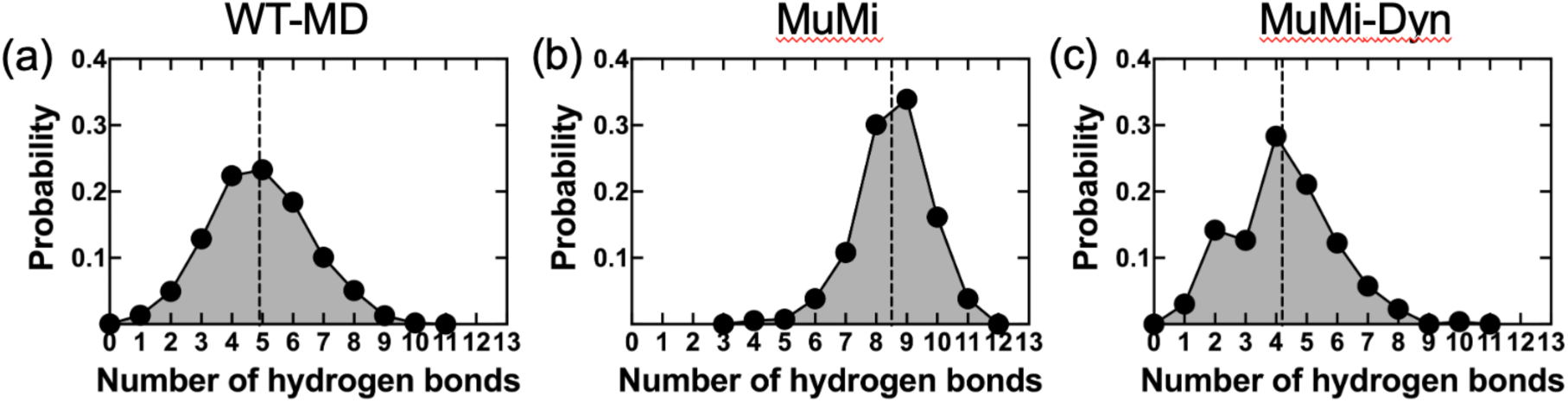
Probability distribution of total hydrogen bonds between GB1 and IgG-Fc for, **(a)** 2 µs-long MD simulations; **(b)** MuMi; and **(c)** MuMi-Dyn. Mean values are indicated by the vertical dashed lines. In minimized structures **(a)**, the number of intermolecular hydrogen bonds varies between 3-12 with an average of 8.5 bonds.

**Figure S6.**
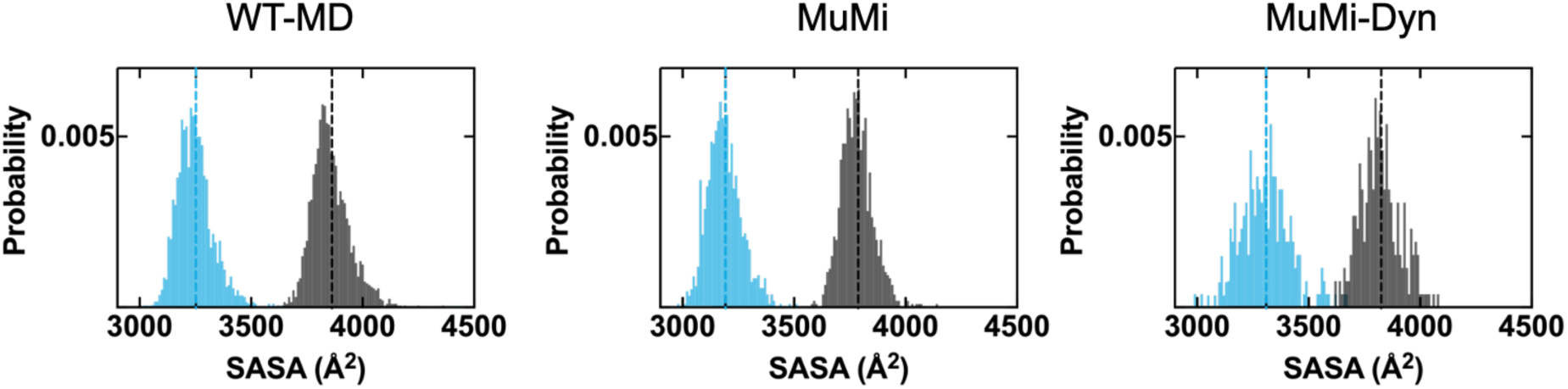
Probability distribution of SASA for unbound (black) and bound (blue) GB1 for, **(a)** 2 µs-long MD simulations; **(b)** MuMi; and **(c)** MuMi-Dyn. Mean values are indicated by the vertical dashed lines. SASA values of minimized WT for unbound and bound forms are 3698 and 3239 Å^2^, respectively. After the 1 ns MD extension of WT form, SASA values increase to 3800 and 3388 Å^2^ in unbound and bound forms, respectively. In bound form, solvent accessibility is decreased for all methods.

**Figure S7.**
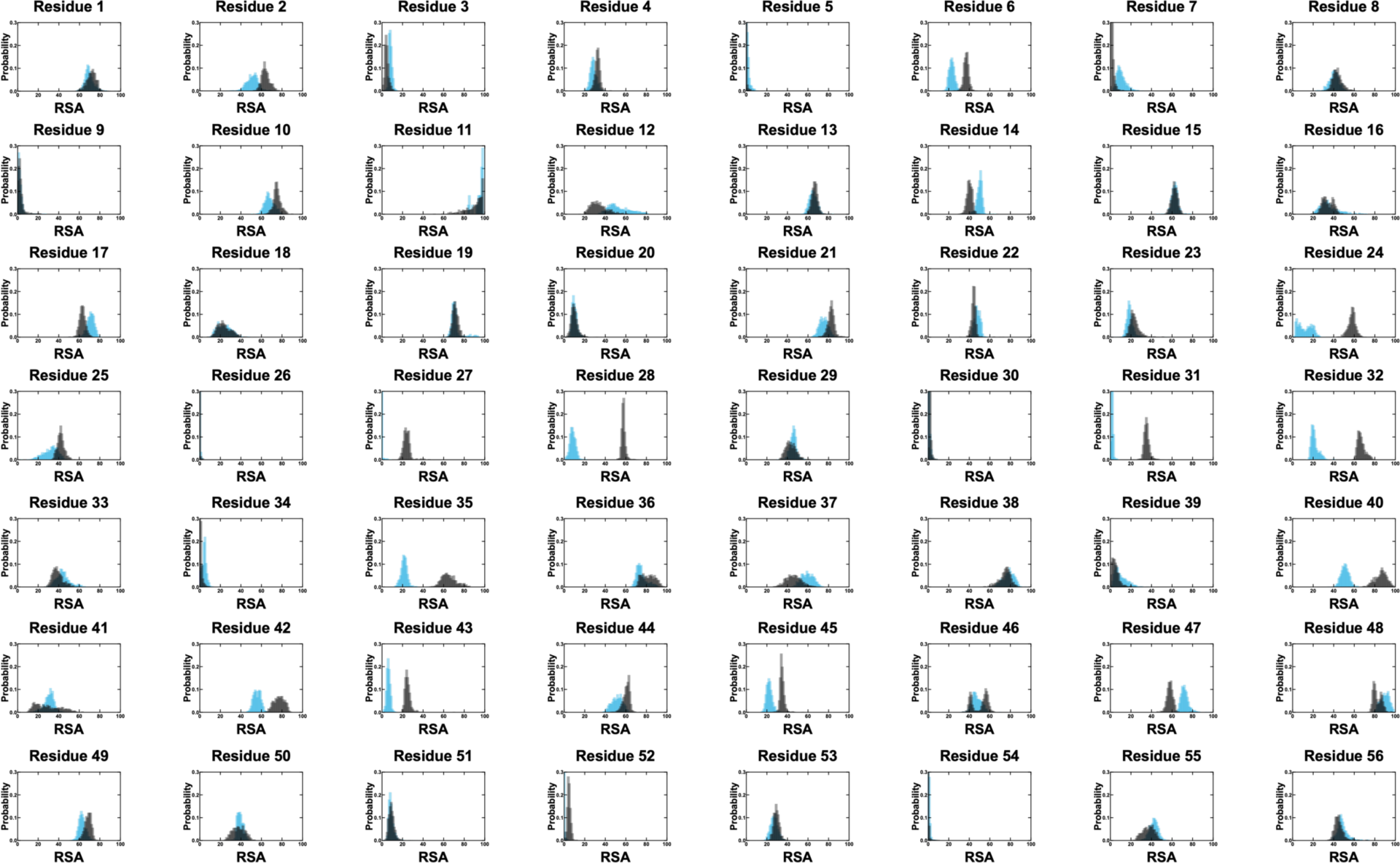
Probability of RSA of unbound (in black) and bound (in blue) GB1 for MuMi. Probabilities are displayed for 56 residue positions.

**Figure S8.**
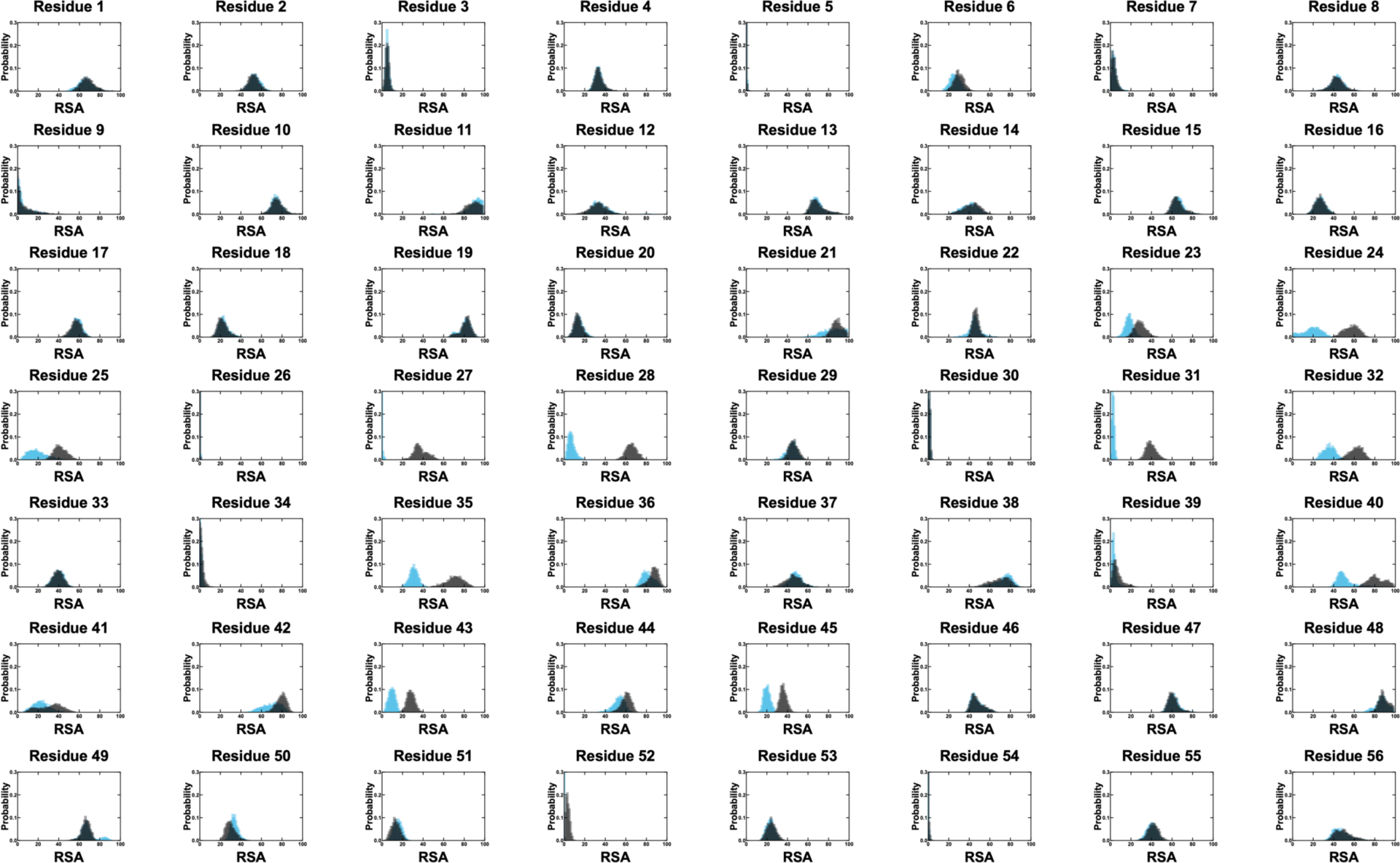
Probability of RSA of unbound (in black) and bound (in blue) for WT MD simulations. Probabilities are displayed for 56 residue positions.

**Figure S9.**
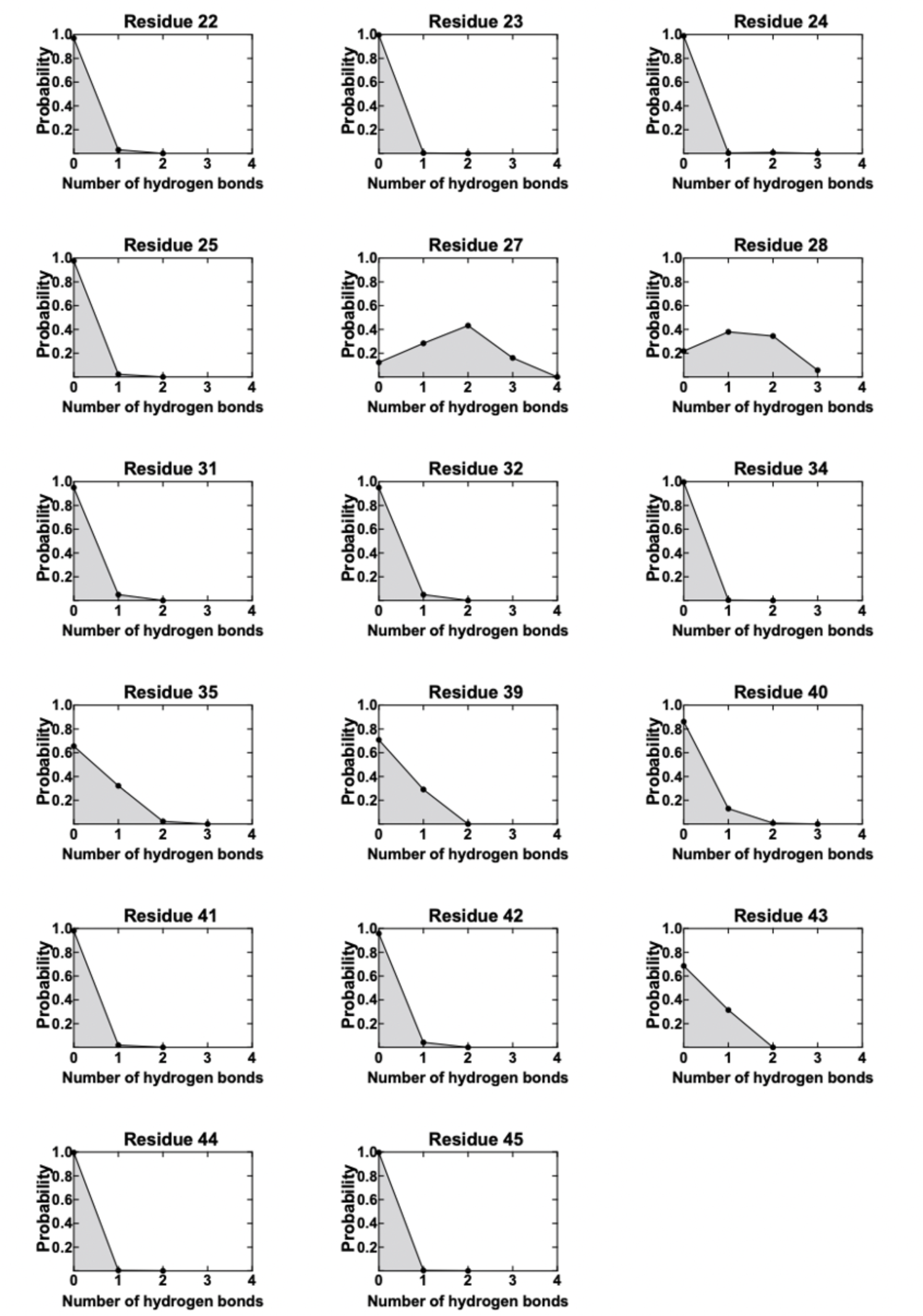
Probability of hydrogen bonds between GB1 and the partner for MuMi-Dyn. All residues having even a single occurrence of hydrogen bonds are shown.

**Figure S10.**
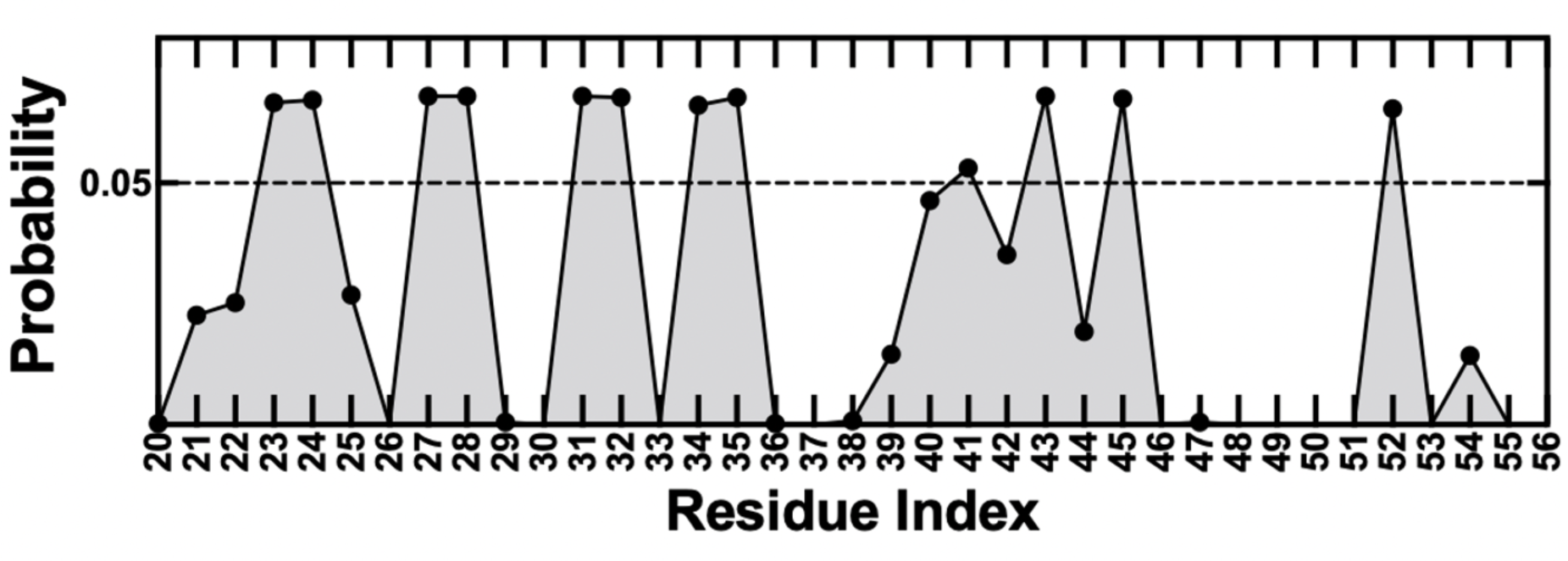
Probability of being located at the binding interface, calculated for MuMi-Dyn. Compared to MuMi and WT MD simulations, residues 40 and 41 have lower probability.

**Figure S11.**
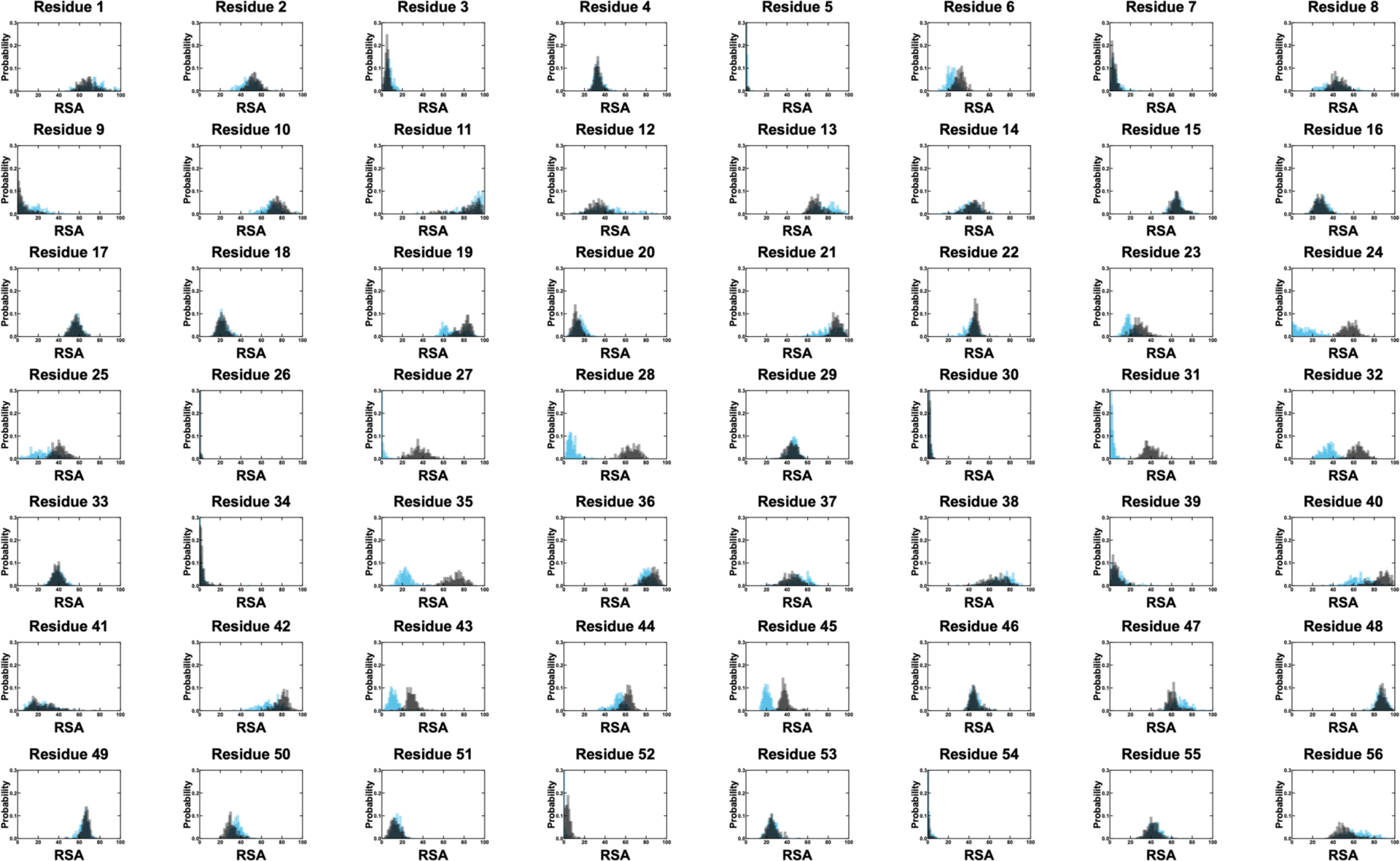
Probability of RSA of unbound (in black) and bound (in blue) for MuMi-Dyn. Probabilities are displayed for the 56 residue positions.

